# Genomic analysis of antimicrobial resistant *Escherichia coli* isolated from manure and manured agricultural grasslands

**DOI:** 10.1101/2024.08.16.608224

**Authors:** C. Tyrrell, C.M. Burgess, F.P. Brennan, D. Münzenmaier, D. Drissner, R.J. Leigh, F. Walsh

## Abstract

Antimicrobial resistance (AMR) is a multifactorial issue involving an intertwining relationship between animals, humans and the environment. The environment can harbour bacteria that are pathogenic to human health, including *Escherichia coli*, an indicator of environmental faecal contamination. Through culture dependent approaches this study identified 46 *E. coli* isolates in porcine and bovine manure, non-manured and manured soil, and the phyllosphere of manured grass. The grass isolation highlights grass as an environmental reservoir for *E. coli.* Whole genome sequencing identified 11 different multi-locus sequence types. We also identified a diverse plasmidome with 23 different plasmid replicon types. The *E. coli* isolates were phenotypically antibiotic resistance, predominantly multidrug resistant. Additionally, whole genome sequencing identified 31 antibiotic resistance genes, and mutations in the *gyrA*, *parC*, and *parE* genes, conferring fluoroquinolone resistance. The main virulence genes were associated actin mediated locomotion (*icsP*/*sopA*), siderophore production and alginate production (*algA*), which suggest adaptation to survive in the non-gut environment or the UV environment of grass surfaces. These results suggest that *E. coli* in soils and grasses may adapt to their new environments evolving novel strategies. This study demonstrates grass as an understudied environmental niche of AMR *E. coli*, which directly links the environment to the grass grazing animal and vice-versa via the circular economy of manure application.

**Impact statement:** *Escherichia coli* is capable of surviving across biomes within One Health. This study sheds light on the genomic elements present in AMR *E. coli* in the understudied niche of agricultural grassland.

**Data summary:** The genome sequences have been deposited in Genbank. Bioproject number PRJNA1080214 and SRP491607 in the sequence read archive https://www.ncbi.nlm.nih.gov/sra/?term=SRP491607.

## Introduction

The One Health concept is critically important to understand the dissemination of antimicrobial resistance (AMR). One Health involves understanding the link between humans, animals and the environment and, in the case of AMR, is particularly relevant due to the ubiquitous nature of antimicrobial resistant bacteria (ARB)^1^. Manure is utilised to enable the circular economy of resources on farms to transfers additional nutrients to grass and soil^2,3^. Furthermore, as these environments yield markedly different conditions to the intestinal tract, evolutionary adaptation can be reasonably expected in isolates observed in these environments^4^. The role of the environment in the spread of AMR and on the occurrence of AMR in environmental bacteria is of interest, particularly in environments associated with human use, such as agricultural land^5–11^. In recent years, the impact manure application has on the occurrence of AMR of agricultural land has received attention due to the AMR selection pressure caused by manure application, and the associated introduction of antimicrobial resistance genes (ARGs), mobile genetic elements (MGEs) and antibiotic residues into the soil^12^. Additionally, bacteria that are clinically important nosocomial pathogens can also be found in the environment, such as the World Health Organisation (WHO) priority pathogens: *Klebsiella pneumoniae, Acinetobacter baumannii, Escherichia coli* and *Pseudomonas aeruginosa.* In terms of antibiotic resistance occurring in these pathogens, carbapenem resistant strains of these species and extended spectrum β-lactamase (ESBL) producing *Enterobacterales* are of critical priority. The plasticity of the *Escherichia coli* genome has been well documented ^2,13^ displaying high amenability to recombination and mobile genetic element uptake ^14–16^. A large degree of this richness can be attributed to the enormous plasmid diversity observed in *E. coli* ^17–20^.

In this study, *E. coli* was isolated and compared from bovine and porcine manure, soil, and grass, demonstrating differential gene profiles on both the chromosome and plasmid sequences. Furthermore, these isolates were compared to a custom database of approximately 27,000 high quality and metadata rich publicly available *E. coli* genomes, demonstrating the adaptability and diversity of this species complex. Using a dual ecological and evolutionary approach for a One Health problem we aimed to characterise antimicrobial resistant *E. coli* in agricultural grassland and to shed light on their potential pathogenicity.

## Methods

### Bacterial isolation from manure, grass, and soil samples

The details of the field trial sampling and sample processing were previously described in detail and do not diverge from this method in this paper^2^. A summary of the sampling is provided for an overview: A grassland field trial was conducted on a research field site in the Southeast of Ireland. Bacterial isolation samples from the following seven timepoints were used: Background (BM) prior to manure application, one week following manure spreading (T1), three weeks following manure spreading (T3), five weeks following manure spreading (T5), ten weeks following manure spreading (T7), 14 weeks following manure spreading (T8) and 18 weeks following manure spreading (T9). Grass samples were prepared for isolation by sonicating 100g of material in 250ml of PBS for 5 minutes using a modified method form (20). Following filtering the liquid through a sterile sieve, 10ml of the sonication liquid was filtered aseptically using a 0.2µm nitrocellulose membrane (Sartorius, Merck). This membrane was then placed into a 50ml falcon tube containing 20ml of nutrient broth (Oxoid) and incubated at 37°C in a shaker at 225rpm (New Brunswick Scientific C25) for 24 hours for bacterial isolation. One gram of manure or soil were added to 20mls of nutrient broth (Oxoid) and incubated for 24 hours at 37°C at 225rpm (New Brunswick Scientific C25). Following the 24-hour enrichment step, the soil and manure samples were left to stand for 5 minutes to allow solid particles to settle. The enriched soil manure and grass samples underwent tenfold serial dilutions (dilution factor = 10^-1^, 10^-2^, 10^-3^, 10^-4^) in sterile PBS. For each enriched manure, grass and soil sample 100μl was used to inoculate Eosin Methylene Blue (EMB) Agar (Oxoid) for the isolation of *E. coli* supplemented with each respective antibiotic at breakpoint concentrations^21^. The antibiotics used were kanamycin (4mg/L), cefotaxime (4mg/L), colistin (4mg/L) and ciprofloxacin (1mg/L). Plates were incubated at 37°C for 24 to 48 hours. Following incubation, presumptive colonies were sub-cultured and purified on low-salt Luria-Bertani (LB) agar (Duchefa) and incubated overnight at 37°C. A maximum of six colonies were picked per agar plate. Glycerol stocks of isolates were prepared and were stored at −80 °C.

### Identification of isolates using MALDI-TOF

A bacterial colony of pure cultures was transferred by direct smearing in duplicate onto spots of the MALDI-TOF mass spectrometry (MS) target (MTP ground steel, Bruker Daltonics) with a tooth-pick. To the dried spots, 1 μL matrix solution (10 mg α-cyano-4-hydroxycinnamic acid, Bruker Daltonics) dissolved in 1 mL acetonitrile-water-trifluoroacetic acid (50:47.5:2.5, (vol/vol/vol), Sigma-Aldrich) was added, and this solution was air-dried. Sample spectra were acquired using a microflex LT MALDI-TOF mass spectrometer (Bruker Daltonics) and the flexControl software *v.*3.4 (Bruker Daltonics). Spectra were classified using the Bruker Taxonomy main spectra database (MBT Compass *v*.4.1. with 8468 spectra present, Bruker Daltonics). Bacterial were identified to the species level if the score value was above 2.00 or to the genus level if the score was between 1.70 and 1.99.

### Antimicrobial susceptibility testing

The *E. coli* (n = 46) underwent antimicrobial susceptibility testing using the EUCAST or CLSI disk diffusion method for susceptibility to cefotaxime (5 µg), imipenem (10 µg), amikacin (30 µg), kanamycin (30 µg), tetracycline (30 µg) and ciprofloxacin (5 µg) (Oxoid). As there are no EUCAST breakpoints for kanamycin and tetracycline CLSI (2020) guidelines and breakpoints were used^21, 22^.

### DNA extraction and whole genome sequencing

Bacterial total DNA was extracted from *E. coli* (n = 46) using the NucleoSpin Microbial DNA Mini kit for DNA from microorganisms (Machery Nagel) according to manufacturer’s instructions. Three *E. coli* samples (39286, 39287, 39288) were sequenced using short read Illumina sequencing by MicrobesNG and the remainder were sequenced by Novogene.

### Dataset construction

All available *Escherichia coli* genome assembly metadata were downloaded from the National Centre for Biotechnology Information using NCBIMeta *v.*0.8.2. As a first filter, genomes with metadata entries for year-of-isolation, isolation source, and country-of-isolation were extracted. The entire list of available genomes were extracted from NCBI Assembly. As a second filter, genomes with fragmented or partial assemblies or with inclusive taxonomies were discarded and definitively genomes labelled as *E. coli* were extracted. Genomes that satisfied the criteria of both filtration steps were downloaded. Each genome was subjected to MLST using mlst *v*.2.19.0 (https://github.com/tseemann/mlst) and genomes not assigned to *E. coli* were discarded. Each genome was separated into chromosomal and plasmid components using PlasClass *v*.0.1.1 with default settings^23^. Each chromosomal genome was annotated using Prokka *v.*1.14.6 with default settings and assessed for completeness using CheckM *v.*1.2.2 under default settings and using the *E. coli* marker gene set^24^. Genomes with a completeness score ≤ 97.5% were discarded. The N50 score for each genome was calculated using Quast *v.*5.2^25^. Chromosomal genomes were used to construct a pairwise similarity network using Mash *v*.2.2.2 with a distance cut-off (*D* ≤ 0.005) and clustered using MCL *v*.14-137 using default settings^26,27^. Chromosomal sequences from isolates sequenced for this study were also used in this clustering step. A representative genome (from publicly available genomes) for each cluster was selected using the longest N50 score. The N50 score was selected as genomes with longer N50 scores provide more reliable pangenome graphs (discussed below). These filtration steps resulted in a comparative dataset of 885 *E. coli* genomes to which our 46 genomes were added, resulting in a total dataset of 931 genomes. These 931 genomes were assessed for population partitioning using PopPunk *v.*2.6.1^28^ with default settings and using the *E. coli* reference dataset provided with the software. This resulted in 859 subpopulations, of which 829 contained a single genome, further demonstrating the genomic diversity of our selected 931 genomes.

### Plasmid partitioning

Each genome sequenced for this study was partitioned into chromosomal and plasmid sequences using platon *v.*1.5.0 using database *v*.1.5 with default settings^29^.

### Pangenome construction

A pangenome graph was constructed from the 931 chromosomal genomes using Roary *v*.3.13.0 using a percentage identity cut-off of 70% (30). This non-default filter was used to reflect the diverse expected pangenome^31^. A second pangenome was constructed using just the chromosomal genomes from isolates sequenced for this study with the same parameters as above. Finally, a third pangenome was constructed using the whole genomes isolates sequenced for this study with the same parameters as above.

### Phylogeny construction

Single copy, ubiquitous gene orthologs were extracted from the pangenome graph (*n =* 805) and aligned using Prank *v*.170427 with default settings^32^. Each alignment was trimmed using TrimAL *v*.1.4rev15 with the “-automated1” flag^33^. All alignments were concatenated into a superalignment (length = 634,861 phylogenetically informative nucleotide sites) using FASconCAT *v*.1.11 under default settings^34^. The superalignment was assessed for the most suitable model of nucleotide evolution using IQTree *v*.2.0.7^35^. The generalised time reversible model with proportional site invariance and gamma distribution models (GTR+I+G) was selected from the 286 sampled models using Aikike’s Information Criterion^36,37^. The superalignment was then used to construct a consensus phylogeny from 10,000 bootstrapped replicates with IQTree2 using the “-B 10000” flag. The resultant phylogeny was visualised using iToL *v.*5^38^. To determine a root for the phylogeny, a second pangenome graph was constructed from the 931 chromosomal genomes as above plus an additional five annotated *Citrobacter fruendii* chromosomal genomes (GenBank IDs = GCA_003665615.1, GCA_003812345.1, GCA_015910405.1, GCA_022646275.1, and GCA_023330605.1). A superalignment was constructed as above from single copy uniquitous genes. A phylogeny was constructed using FastTree *v*.2.1 using the “-nt” (nucleotide specific) flag and the GTR+I+G model^39^. This phylogeny was rooted at the branch representing the divergence of *C. fruendii* from *E. coli.* The earliest branching *E. coli* lineage according to this phylogeny was selected as the root of the 931-genome phylogeny. This rooting strategy was confirmed using FastRoot (MinVar) *v*.1.5 under default settings (https://github.com/uym2/MinVar-Rooting). Finally, to allow greater exploration into isolates sequenced for this analysis, a pangenome graph was constructed for these 46 genomes using Roary and each gene family was aligned and trimmed using Prank and TrimAl as above; ubiquitous, single copy gene alignments (*n* = 3,386) were extracted and concatenated into a superalignment (length = 3,251,889 bp). Again, this superalignment was used to construct a consensus phylogeny with 10,000 bootstrap replicates using IQTree2 as above, where the GTR+I+R4 model was selected as the most appropriate model of nucleotide evolution. The root, as selected by FastRoot, was determined to be Isolate 10, in agreement with the other two phylogenies. The phylogeny was visualised using iToL as above.

### Phylogenetic clustering

The 931-genome phylogeny was partitioned into phylogenetic groups using FastBAPS *v*.1.08 using the super alignment as support^40^. These results were used to annotate the previously constructed phylogeny.

### Genome annotation

Each chromosomal genome was annotated for antimicrobial resistance using (https://github.com/tseemann/abricate) with the comprehensive antimicrobial resistance database for biocide and heavy metal resistance using ABRicate with a backtranslated version of BacMet *v*2.0, for virulence factors using the virulence factor database (VFDB), for O- and H-antigen types using the EcOH *v*.2 dataset^41^. The CARD, VFDB, and EcOH, databases are provided as standard with ABRicate and the backtranslated version of BacMet was obtained from a clinical microbiology study on vancomycin resistant *Enterococcus faecium*^42^. Resistance genotypes associated with point mutations were derived using PointFinder *v*.3.1.1. with the *E. coli* reference dataset provided with PointFinder^43^. Previously assigned MLSTs were used to assign genomes to clonal complexes with PubMLST^44^. Additionally, each chromosome and plasmid sequence from isolates sequenced for this study were annotated using Bakta *v.*1.5.1 using database *v.*1.5^45^. This secondary analysis was performed to restrict the ambiguity arising from hypothetical proteins as assigned by Prokka. Plasmid sequence typing was performed on isolates sequenced for this study using ABRicate with the PlasmidFinder database^46^.

## Results

*Escherichia coli* were isolated only from manure treated grass (n = 14) (i.e. no grass without manure contained *E. coli* across timepoints 1, 3, 5 and 10 from bovine, porcine or poultry manured grass. The *E. coli* were isolated from bovine and porcine manure (n = 21), but not poultry manure (Table 1). They were also isolated from soil across timepoints 1, 3 and 10, to which no manure was added (n = 9) or porcine manure was added (n = 2).

**Table 1.** Cluster and sequence types of all isolates.

| *E. coli*<br>isolate | OH group | Sequence<br>type | Clonal cluster | Phylogroup | Chromosome<br>cluster | Chromosome<br>group | Plasmid group | Set | Timepoint |
| --- | --- | --- | --- | --- | --- | --- | --- | --- | --- |
| 1 | O8H25 | 58 | ST155 complex | ξ | Cluster_9 | CG1 | PG1B | Porcine | pre application |
| 10 | O17H18 | 106 | ST69 complex | γ | Cluster_6 | CSG1 | PG1C | Bovine | pre application |
| 11 | O17H25 | 1126 | ST155 complex | ξ | Cluster_9 | CG1 | PG1C | Bovine | pre application |
| 12 | O8H25 | 1126 | ST155 complex | ξ | Cluster_9 | CG1 | PG1C | Bovine | pre application |
| 13 | O8H25 | 58 | ST155 complex | ξ | Cluster_9 | CG1 | PG1B | Bovine | pre application |
| 14 | O8H30 | 1431 | NaN | ξ | Cluster_17 | CG2 | PG2 | Bovine | pre application |
| 15 | O8H30 | 1431 | NaN | ξ | Cluster_17 | CG2 | PSG1 | Bovine | pre application |
| 16 | O8H30 | 1431 | NaN | ξ | Cluster_17 | CG2 | PG2 | Bovine | pre application |
| 17 | O8H30 | 1431 | NaN | ξ | Cluster_17 | CG2 | PG2 | Bovine | pre application |
| 18 | O8H30 | 1431 | NaN | ξ | Cluster_17 | CG2 | PG2 | Bovine | pre application |
| 19 | O8H25 | 1126 | ST155 complex | ξ | Cluster_9 | CG1 | PG1C | Bovine | pre application |
| 2 | O8H26 | 156 | ST156 complex | ξ | Cluster_32 | CG3 | PG3 | Porcine | pre application |
| 20 | O8H25 | 58 | ST155 complex | ξ | Cluster_9 | CG1 | PG1B | Soil | Week 1 |
| 21 | O8H25 | 58 | ST155 complex | ξ | Cluster_9 | CG1 | PG1B | Soil | Week 1 |
| 22 | O8H25 | 58 | ST155 complex | ξ | Cluster_9 | CG1 | PG1B | Soil | Week 1 |
| 23 | O8H25 | 58 | ST155 complex | ξ | Cluster_9 | CG1 | PG1B | Soil | Week 1 |
| 24 | O8H25 | 58 | ST155 complex | ξ | Cluster_9 | CG1 | PG1B | Soil | Week 1 |
| 25 | O8H25 | 58 | ST155 complex | ξ | Cluster_9 | CG1 | PG1B | Soil | Week 1 |
| 26 | O88H8 | 446 | ST446 complex | ξ | Cluster_77 | CG4 | PG4 | Soil | Week 10 |
| 27 | O88H8 | 446 | ST446 complex | ξ | Cluster_77 | CG4 | PG4 | Soil | Week 10 |
| 28 | O88H8 | 446 | ST446 complex | ξ | Cluster_77 | CG4 | PG4 | Soil | Week 10 |
| 29 | O88H8 | 446 | ST446 complex | ξ | Cluster_77 | CG4 | PG4 | Soil | Week 10 |
| 3 | none-H26 | 156 | ST156 complex | ξ | Cluster_32 | CG3 | PG3 | Porcine | pre application |
| 30 | O163H6 | 189 | ST165 complex | v | Cluster_36 | CG6 | PG6 | Grass | Week 1 |
| 31 | O100H32 | 10 | ST10 complex | v | Cluster_450 | CG7 | PG7 | Grass | Week 1 |
| 32 | O100H32 | 10 | ST10 complex | v | Cluster_450 | CG7 | PG7 | Grass | Week 1 |
| 33 | O163H6 | 189 | ST165 complex | v | Cluster_36 | CG6 | PG6 | Grass | Week 1 |
| 34 | O23H16 | 453 | ST86 complex | ξ | Cluster_24 | CSG2 | PSG4 | Grass | Week 3 |
| 35 | O8H25 | 58 | ST155 complex | ξ | Cluster_9 | CG1 | PG1A | Grass | Week 5 |
| 36 | O8H25 | 58 | ST155 complex | ξ | Cluster_9 | CG1 | PG1A | Grass | Week 5 |
| 37 | O8H25 | 58 | ST155 complex | ξ | Cluster_9 | CG1 | PG1A | Grass | Week 5 |
| 38 | O8H25 | 58 | ST155 complex | ξ | Cluster_9 | CG1 | PG1A | Grass | Week 5 |
| 39 | O8H25 | 58 | ST155 complex | ξ | Cluster_9 | CG1 | PG1A | Grass | Week 5 |
| 39286 | O187H52 | 1248 | NaN | ξ | Cluster_378 | CSG3 | PSG5 | Soil | Week 3 |
| 39287 | O187H10 | 617 | ST10 complex | v | Cluster_23 | CG5 | PSG2 | Porcine | pre application |
| 39288 | Onovel32H26 | 156 | ST156 complex | ξ | Cluster_32 | CG3 | PG3 | Porcine | pre application |
| 4 | none-H26 | 156 | ST156 complex | ξ | Cluster_32 | CG3 | PG3 | Porcine | pre application |
| 40 | O8H25 | 58 | ST155 complex | ξ | Cluster_9 | CG1 | PG1A | Grass | Week 5 |
| 41 | O8H25 | 58 | ST155 complex | ξ | Cluster_9 | CG1 | PG1A | Grass | Week 5 |
| 42 | O8H25 | 58 | ST155 complex | ξ | Cluster_9 | CG1 | PG1A | Grass | Week 5 |
| 43 | O8H25 | 58 | ST155 complex | ξ | Cluster_9 | CG1 | PG1A | Grass | Week 5 |
| 5 | Onovel32H10 | 617 | ST10 complex | v | Cluster_23 | CG5 | PSG3 | Porcine | pre application |
| 6 | O8H30 | 1431 | NaN | ξ | Cluster_17 | CG2 | PG2 | Bovine | pre application |
| 7 | O8H30 | 1431 | NaN | ξ | Cluster_17 | CG2 | PG2 | Bovine | pre application |
| 8 | O8H30 | 1431 | NaN | ξ | Cluster_17 | CG2 | PG2 | Bovine | pre application |
| 9 | O8H30 | 1431 | NaN | ξ | Cluster_17 | CG2 | PG2 | Bovine | pre application |

NaN = no ST complex assigned.

The antimicrobial resistance profiles of the *E. coli* (n = 46) comprised, 35 tetracycline resistant (n = 35), 9 were cefotaxime resistant (n = 9), 24 were kanamycin resistant (n = 24), or ciprofloxacin resistant (n = 16). There were no amikacin or imipenem resistance. Multi-drug resistance was present in 31 isolates, with 18 conferring resistance to two antimicrobial classes and 13 resistant to three antimicrobial classes. *Escherichia coli* isolates displaying resistance to two classes were resistant to tetracycline and kanamycin or tetracycline and ciprofloxacin. Isolates demonstrating resistance to three classes of antibiotics were resistant to tetracycline, kanamycin and ciprofloxacin or tetracycline, cefotaxime and ciprofloxacin. The cefotaxime resistant *E. coli* were isolated from bovine manure and tested negative for AmpC β-lactamase production and positive for extended spectrum beta-lactamase (ESBL) production (Supplementary information).

The bacterial genome data from our isolated *E. coli* was clustered to identify patterns between our isolate genomes and all available genomes of *E. coli*. The complete dataset (our isolates and all available *Escherichia coli* genome assembly metadata were downloaded from the National Centre for Biotechnology Information using NCBIMeta *v.*0.8.2 that passed the filter criteria) was collapsed to 931 genomes (885 genomes from public sources and the 46 isolated for this study). Of the 885 genomes from public sources, 579 were from groups with *n* ≥ 2 genomes and 306 were singletons (Supplementary Information (SI) Table Cluster). Groups ranged from two to 4,458 genomes with a highly skewed distribution; most groups (*n* = 509; 87.91%) were in groups *n* ≤ 50 (µ = 46.56±235.6)*, Q*_0.25_ = 3; *Q*_0.5_ = 5; *Q*_0.75_ = 17). Isolates sequenced for this study were observed in ten clusters (6, 9, 17, 23, 24, 32, 36, 77, 378, 450) (Table 1). As our isolates were of recent origin we aimed to identify the earliest isolate from each cluster to investigate whether these isolates were recent emerging isolates or have been detected previously, their range of global distribution and if the cluster isolates were from a specific sample type e.g. agriculture only or human sources.

Cluster 6 was composed of 862 genomes, the earliest of which was isolated from a human haematological sample in 1983, USA, based on the sample metadata. Since 1983, cluster 6 has also been observed in 53 countries, spanning six continents. The majority (*n* = 743; 86.2%) of cluster 6 isolates were from human isolation. The remaining isolates were of animal origin including bovine and porcine samples. Most isolates were of gastrointestinal or faecal origin. From a food safety perspective, cluster 6 isolates have been observed in bovine milk and poultry egg products.

Cluster 9 comprised 728 genomes, the oldest of which was sampled from bovine and caprine gastrointestinal and immunological samples in 1970. The first cluster 9 human isolate was observed in a Canadian urinary tract infection sampled in 1980. Since then, cluster 9 has been isolated in 55 countries across six continents. Since its emergence, cluster 9 has been observed across a multitude of taxa (mammals, birds, and molluscs) from wild animal, veterinary, agricultural and clinical samples. Cluster 9 has also been observed in a plethora of clinical tissue types in infected humans and animals, and has also been observed in meat, milk and egg samples.

Cluster 17 was composed of 304 genomes with the oldest isolate being isolated from a goat immunological sample in 1971, USA. Since 1971, cluster 17 has been observed in 41 countries, spanning six continents. Most isolates were of human origin (*n* = 141; 46.4%). The remaining animal samples were across a range of animals including of bovine, poultry, and porcine and were mostly of gastrointestinal or faecal origin. From a food safety perspective, cluster 17 was also observed in bovine milk and poultry egg samples.

Cluster 23 contains 211 genomes with the earliest sample isolated from a human urinary tract infection in France, 2002. Since 2002, cluster 23 has been isolated in 37 countries, spanning six continents. The majority (*n* = 185; 87.7%) of cluster 23 genomes were isolated from human clinical samples. Cluster 23 has also been isolated from animals including poultry, bovine, and porcine origins. Most non-human isolates were observed in gastrointestinal or faecal samples.

Cluster 24 comprised 206 genomes, with the oldest isolate from a porcine respiratory sample in 1975. The first cluster 24 human isolate was observed in a gastrointestinal tract sample from Chile in 1986. Since 1975, cluster 24 has been observed in 37 countries, across six continents.

Cluster 24 has been observed in wild mammals and birds, domestic and agricultural mammals and birds, and in human samples. From a food safety perspective, cluster 24 was also isolated from duck egg, pork, and bovine dairy samples.

Cluster 32 was composed of 131 genomes with the earliest being isolated from wild deer in 1981, USA. Since 1981, cluster 32 has been observed in 33 countries, spanning six continents. The majority of cluster 32 isolates were of human isolation (*n* = 78; 59.5%), followed by poultry samples (*n* = 27; 20.6%), and bovine samples (*n* = 7; 5.3%). Regarding food safety, cluster 32 was also observed in one beef, two milk and one poultry meat sample.

Cluster 36 was composed of 108 genomes with the oldest isolate being sampled from a 1979 poultry faecal sample in the USA. The first cluster 36 human isolated was sampled from a Saudi Arabian gastrointestinal sample from 1991. Since 1979, cluster 36 has been observed in 23 countries spanning six continents. In addition to humans and poultry, cluster 36 has been observed in agricultural (duck, bovine, and porcine samples) and wild animals (mustelids and pigeons). Within agricultural animals, cluster 36 was observed in gastrointestinal and faecal samples for the three species, within poultry cardiac and hepatological samples, and within porcine hepatological obstetric, and respiratory samples. From a food safety perspective, cluster 36 was also observed in milk samples. 6, 9, 17, 23, 24, 32, 36, 77, 378, 450

Cluster 77 was composed of 45 genomes with the earliest isolated sample being isolated from a bovine faecal sample from 1988, USA. The majority of isolates are of human origin, with almost all from gastrointestinal or faecal samples.

Cluster 378 was composed of three isolates sourced in the USA and one isolate from this study. Two isolates were from bovine gastrointestinal samples (observed in 2016) and one human urinary tract sample observed in 2020.

Cluster 450 was composed of seven isolates, including two from this study. One isolate was observed in a UK poultry sample (observed in 2014) and four from human gastrointestinal or urinary samples from China, USA and United Arab Emirates.

### Determination of clonal genomes

Isolates sequenced for this study that were clustered into the same genome cluster and shared a clonal complex were considered to be clonal^44^. With the exception of clusters 9 and 77, all other isolates in the same cluster were sampled from the same source in this study; this may be due the relatively small subset sizes (n = 1 to 4), with the exception of cluster 17 (*n* = 9). Cluster 77 isolates were sampled from week 10 and comprised control soil i.e. no manure added and soil with porcine manure added. Cluster 9 isolates were isolated from bovine manure, bovine manure treated grass, porcine manure, and control soil. The isolates were isolated in both bovine and porcine manure pre-field application and the control soil in week 1. In addition, at week five they were isolated from grass with bovine manure applied but not the porcine manured grass. There were also no isolates from the control grass nor earlier grass samples. The sequence type of isolates from each grouping was ST58 and the plasmid group comprised two groups, suggesting that these are very highly related isolates. This suggests that the isolates on the grass originated from either the soil or the bovine manure and required four weeks to detectable by culture from grass but were not maintained on grass in the subsequent weeks.

For ease of reading, clusters shall be referred to as “chromosomal groups” (CG) with the following reassignments (based on the size of a given CG): cluster 9 = CG1, cluster 17 = CG2, cluster 32 = CG3, cluster 77 = CG4, cluster 23 = CG5, cluster 36 = CG6, cluster 450 = CG7; as clusters 6, 24, and 378 only had one associated isolate sequenced for this study, these are referred to “chromosome singleton groups” (CSG) and assigned to CSG1, CSG2, and CSG3 respectively.

When plasmid derived sequences from this study were clustered, eight distinct plasmid groups (PG) were observed (Table 1; Figure 1). Plasmid groups largely followed chromosomal groups with the exception of CG1. For CG1, isolates were distributed between PG 1A, 1B or 1C with all CG1 bovine manure treated grass assigned to in PG1A (*n* = 9), all control soil cluster 9 isolates, the single porcine manure isolate, and one bovine manure isolate assigned to PG1B (*n* = 8), and the remaining bovine manure treated isolates were assigned to PG1C (*n* = 3). For CG2 (all bovine manure isolates), eight of nine isolates were assigned to PG2 (*n* = 8) and one isolate presented a unique plasmid set (PSG1). All CG3 isolates (all porcine manure isolates) were assigned to PG3 (*n* = 4). All CG4 isolates (three control soil isolates and one porcine treated soil isolate) were assigned to PG4 (*n* = 4). Both CG5 isolates (porcine manure isolates) displayed unique plasmid sets (PSG2 and PSG3). CG6 and CG7 isolates (all porcine manure treated grass isolates) were assigned to PG7 and PG8, respectively. The individual isolates in CGs 6, 24 and 378 contained unique plasmids assigned to PSG6, PSG4 and PSG5, respectively.

**Figure 1:**
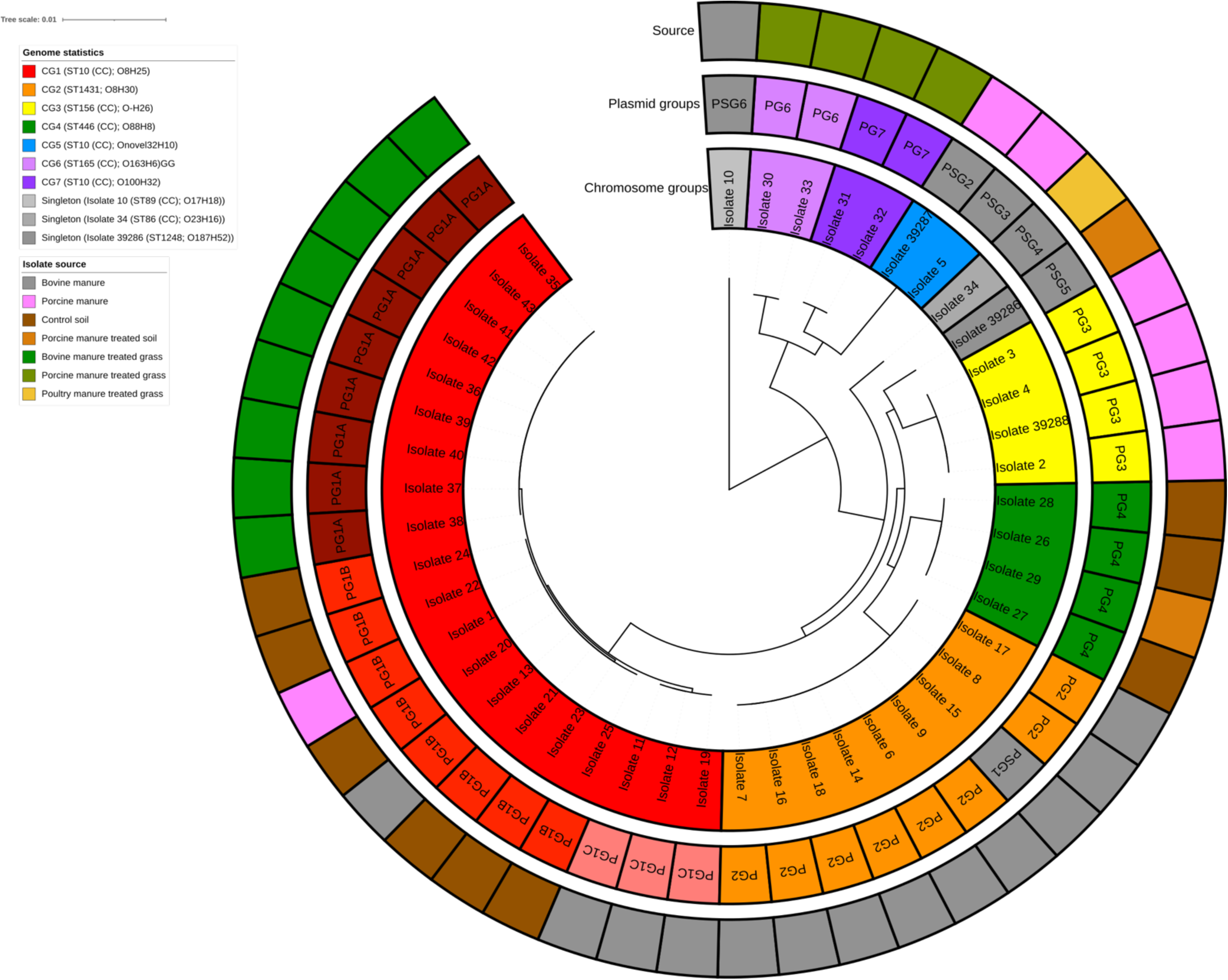
Chromosome and plasmid phylogeny of manure and manured agricultural grasslands *Escherichia coli*

### Multilocus sequence typing of isolates sequenced for this study

A total of 11 sequence types (STs) were observed across isolates sequenced for this study (Table 1, SI Table Genomics). These STs could be further categorised to seven clonal complexes (CC). Neither ST1248 nor 1431 could be assigned to a CC. All CG1 isolates belonged to CC-ST155, with all isolates possessing PG1A and PG1B belonging to ST58 and PG1C belonging to ST1126. All other CGs returned a single ST per group.

### O-antigen H-fimbriae typing

A total of eight O-antigens (O8, O17, O23, O_novel_32, O88, O100, O163, and O187; two isolates (isolates 3 and 4; CG3) did not possess an O-antigen) and ten H-fimbriae (H6, H8, H10, H16, H18, H25, H26, H30, H32, and H52) were observed across the 46 sequenced isolates, culminating to 14 serotypes (Table 1, SI Table Genomics). With the exception of isolate 11, all CG1 isolates presented the O8:H25 serotype, isolate 11, by contrast, presented the O17:H25 serotype. All CG2 isolates were of serotype O8:H30. All CG3 isolates possessed H26 but three different O-antigen combinations were present resulting in three serotypes (H26, O_novel_32:H26, and O8:H26. All CG4 isolates were of serotype O88:H8. Both CG5 isolates had H10 fimbriae but different O-antigens. Both CG6 isolates contained the O163:H6 serotype and both CG7 isolates presented the O100:H32 serotype. The serotypes for CSG1, CSG2, and CSG3 were each unique.

### Plasmid typing of isolates sequenced for this study

When plasmid families were considered, all plasmid sets within each PG (except PG1A, PG1B, and PG7) displayed the same plasmid replicon types (SI Table PlasmidFinder). All PG1A contained Col440II and IncFIB, however only five isolates contained Col156. All PG1B isolates possessed Col8282 and IncFIB, and all but one (isolate 24) possessed ColpVC, six of eight isolates possessed Col(BS512) and four isolates possessed Col156. One isolate contained the IncFIB replicon and the other within the PG7 group did not. Both contained the IncN and Col440II replicons. All PG1C isolates possessed Col440II or ColRNAI and IncFIB. All PG2 isolates possessed IncY only. All PG3 isolates possessed IncFII and p0111. All PG4 isolates possessed IncFIB only. Both PG6 isolates contained IncX2 and IncX5. The PSG1 and PGS2 contained no replicon gene. The remaining PSG plasmids were all IncFIB and PSG5 also contained Col440II_1.

### Phylogeny

The single copy, ubiquitous gene phylogeny of all isolates analysed split into 14 phylogroups (Figure 2). Phylogroups were annotated in descending order based on their divergence from the root. The phylogroups numbers varied from α which was composed of one isolate to ξ which was composed of 297 isolates. The isolates from this study were contained in three phylogroups (Figure 2). Most were from the latest diverging phylogroups ν (*n* = 6) and ξ (*n* = 39) and one isolate in phylogroup γ.

**Figure 2:**
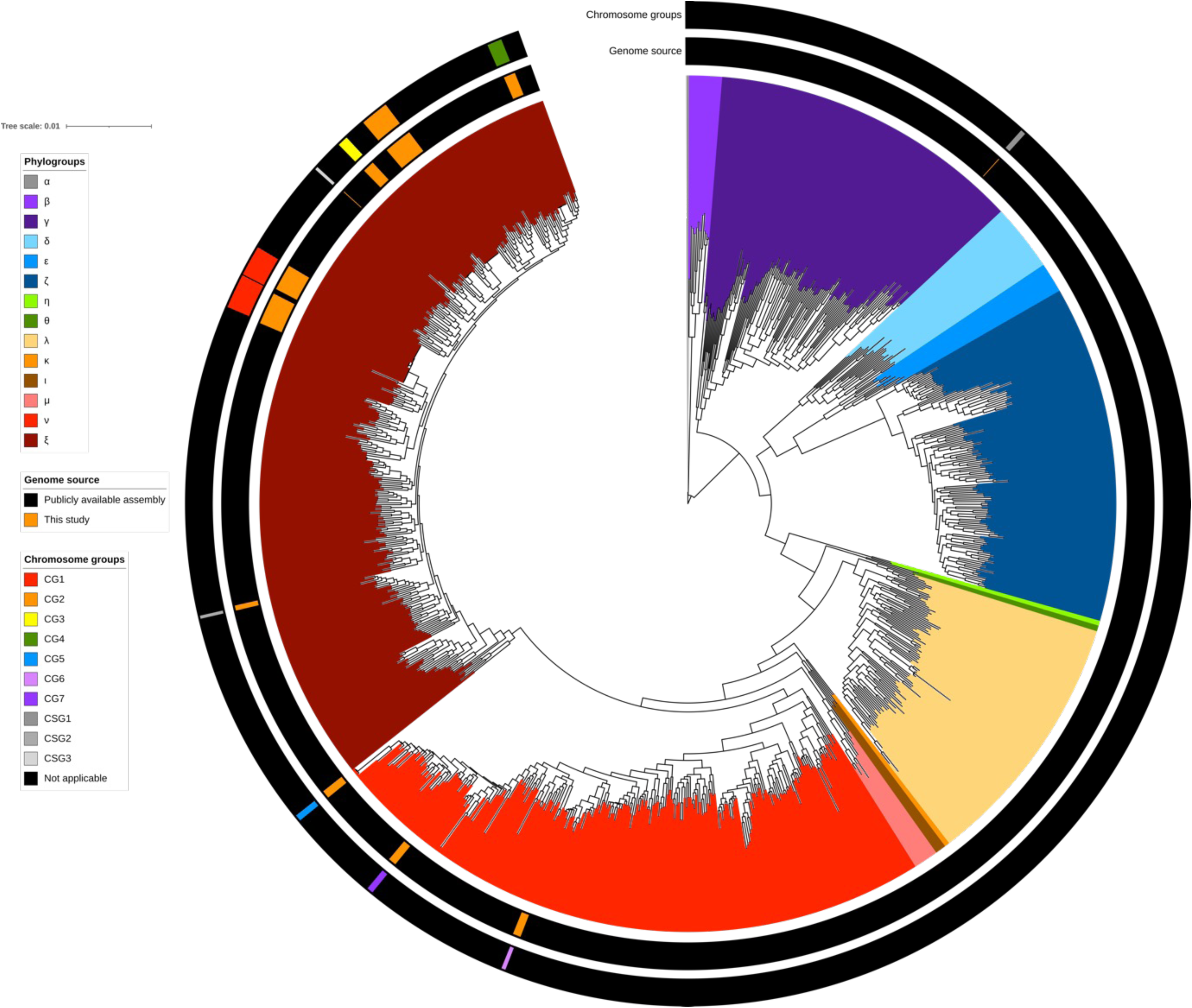
Phylogeny of all *E. coli* isolates analysed in this study The branch length for phylogroup was excessively long and visually obscured all other phylogenetic branches. To counter this, this branch was divided by 100 to allow for better visualisation. The group α serves as an outgroup.

### Pangenomics

When all chromosomal genomes were considered, 46,546 genes were observed (Figure 3). Of these 1,761 were core genes (of which 805 were ubiquitously distributed), 555 were soft core genes, 3,195 were shell genes, and 41,035 were cloud genes (of which 15,478 were singletons and 4,975 were doubletons). When just chromosomal genomes of isolates sequenced for this study were considered, 7,811 genes were observed. Of these, 3,381 were core genes (all of which were ubiquitous), 242 were soft core genes, 1,673 were shell genes, and 2,515 were cloud genes (of which, 1,122 were singletons and 611 were doubletons). When the entire genomes of isolates sequenced for this study (chromosomal and plasmid sequences) were considered, 8,756 genes were observed. Of these, 3,381 were core (again, all of which were ubiquitous), 239 were soft core genes, 1,984 were shell genes, and 3,152 were cloud genes (of which 1,168 were singletons and 906 were doubletons).

**Figure 3.**
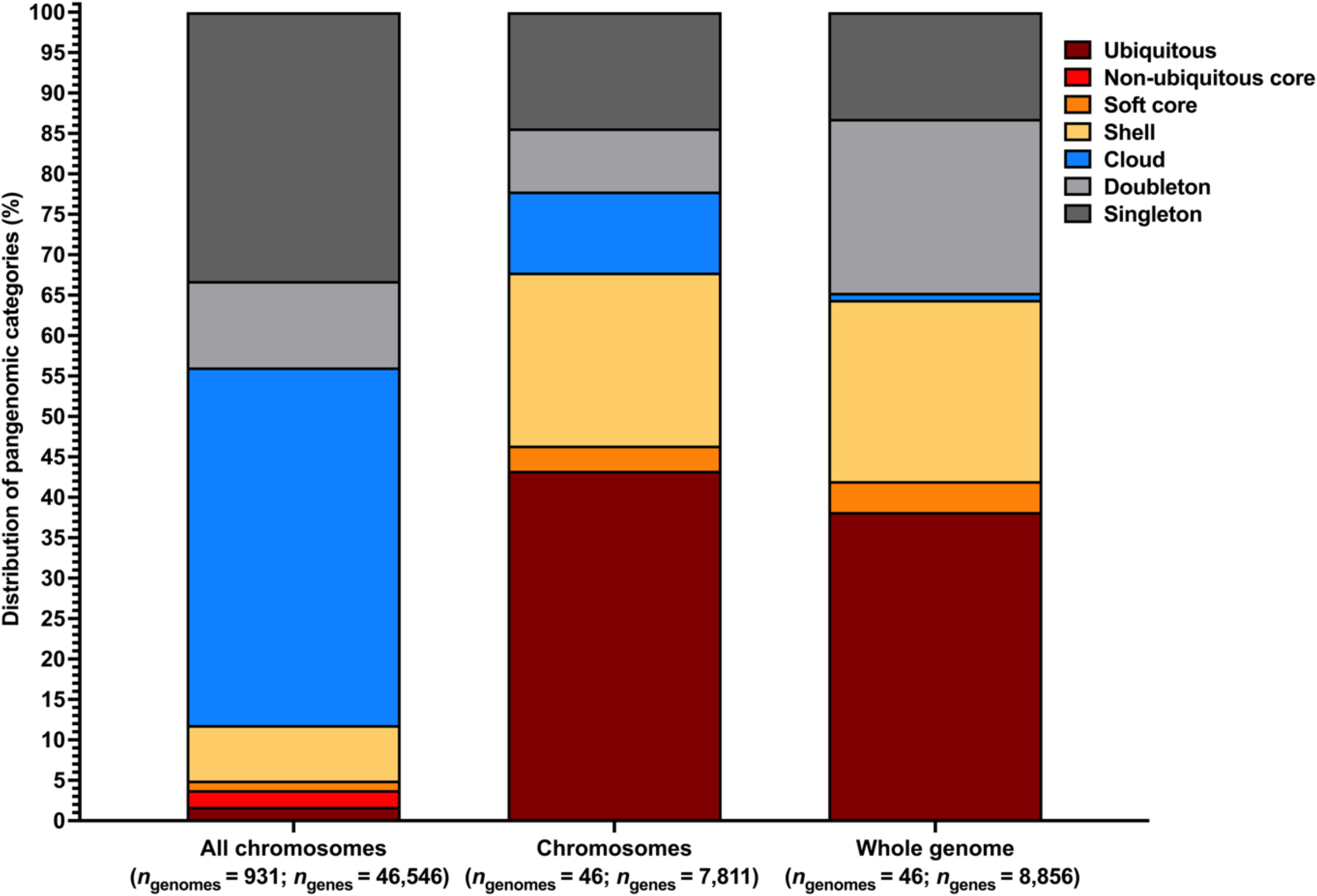
Pangenomic proportions for each genome set The first column represents the pangenomic diversity across the *E. coli* chromosomal phylogeny. The second column illustrates the chromosomal pangenomic diversity within isolates sequenced from this study. The final column illustrates the total pangenomic diversity (chromosomal and plasmid sequences) of isolates sequenced from this study.

### Antimicrobial, metal and biocide resistance and virulence genes

All isolates were susceptible to amikacin and all, except isolate 36, were susceptible to imipenem (10 μg) with isolate 36 displaying intermediate resistance (Table 2). Ciprofloxacin resistance was phylogenetically distributed (associated with chromosomal groups) irrespective of plasmid groups, with all CG2, CG3, CG5, and isolate 34 (CSG2) displaying resistance. All CG2 isolates displayed cefotaxime resistance. Tetracycline resistance was observed in 35 isolates, with susceptibility observed in CG4 isolates, four CG1 isolates, one CG3 isolate was intermediate resistant and the CSG1 and CSG3 isolates also susceptible. Resistance to kanamycin was observed in all CG1 isolates except isolate 36, all CG3 isolates, and isolate 10.

**Table 2.** Antimicrobial resistance phenotypes of all isolates. S = susceptible, I = intermediate, R = resistant.

| Isolates | Tetracycline | Cefotaxime | Kanamycin | Amikacin | Ciprofloxacin | Imipenem |
| --- | --- | --- | --- | --- | --- | --- |
| 1 | R | S | R | S | S | S |
| 2 | R | S | R | S | R | S |
| 3 | R | S | R | S | R | S |
| 4 | R | S | R | S | R | S |
| 5 | R | S | S | S | R | S |
| 6 | R | R | S | S | R | S |
| 7 | R | R | S | S | R | S |
| 8 | R | R | S | S | R | S |
| 9 | R | R | S | S | R | S |
| 10 | S | S | R | S | S | S |
| 11 | S | S | R | S | S | S |
| 12 | S | S | R | S | S | S |
| 13 | R | S | R | S | S | S |
| 14 | R | R | S | S | R | S |
| 15 | R | R | S | S | R | S |
| 16 | R | R | S | S | R | S |
| 17 | R | R | S | S | R | S |
| 18 | R | R | S | S | R | S |
| 19 | S | S | R | S | S | S |
| 20 | R | S | R | S | S | S |
| 21 | R | S | R | S | S | S |
| 22 | R | S | R | S | S | S |
| 23 | R | S | R | S | S | S |
| 24 | R | S | R | S | S | S |
| 25 | R | S | R | S | S | S |
| 26 | S | S | S | S | S | S |
| 27 | S | S | S | S | S | S |
| 28 | S | S | S | S | S | S |
| 29 | S | S | S | S | S | S |
| 30 | R | S | S | S | S | S |
| 31 | R | S | S | S | S | S |
| 32 | R | S | S | S | S | S |
| 33 | R | S | S | S | S | S |
| 34 | R | S | S | S | R | S |
| 35 | R | S | R | S | S | S |
| 36 | S | S | S | S | S | I |
| 37 | R | S | R | S | S | S |
| 38 | R | S | R | S | S | S |
| 39 | R | S | R | S | S | S |
| 40 | R | S | R | S | S | S |
| 41 | R | S | R | S | S | S |
| 42 | R | S | R | S | S | S |
| 43 | R | S | R | S | S | S |
| 39286 | S | S | S | S | S | S |
| 39287 | R | S | S | S | R | S |
| 39288 | I | S | R | S | R | S |

A total of 64 antimicrobial resistance genes were observed across all chromosomal components sequenced for this study (SI Table CARD: Chromosome). However, the vast majority of these genes are efflux genes, which must be actively upregulated to provide the associated resistance phenotype or must the gene is a chromosome gene which must contain a mutation to confer resistance. Therefore, we have highlighted in red the genes that confer resistance by their presence. All PG1A and all CG3 isolates possessed *tetB* and *bla*_TEM-1_ which confer resistance to tetracycline and beta-lactams, respectively. A total of five CG1 isolates possessing PG1B (1, 20, 23-25) possessed *bla*_TEM-190_, an ESBL gene. Both CG6 isolates possessed *ANT(3’’)-IIa* and *SAT-2* which confer resistance to aminoglycosides and nucleosides, respectively. Isolate 39288 contained resistance genes, *sul1*, *AAC(6’)-Ib7*, *catB3*, *catI*, *dfrA1*, which, confer resistance to sulphonamides, aminoglycosides, phenicols (*catB3* and *catI*), and trimethoprims. Finally only isolate 34 contained *bla_TEM-150_*, an ESBL producing gene, on the chromosome.

With the exception of PG4 and isolates in PSG1 and PSG5, all isolates possessed plasmid mediated resistance genes (SI Table CARD: Plasmid). Three efflux genes were observed on plasmids, *floR*, *tet(A),* and *tet(B)*, which confer resistance to phenicols and tetracyclines, respectively. From a distribution perspective, *tet(A)* and *tet(B)* were mutually exclusive, with *tet(A)* contained within all PG1B, PG2, PG6, PG7 and PSG4 isolates (*n* = 20). By contrast, *tet(B)* was only observed in PSG1 and PSG2 (both CG5) isolates. All PG1B, PG1C, and PSG6 isolates possessed *floR*. When non-efflux mediated resistance genes were considered, *APH(6)-Id* (aminoglycoside resistance) was observed in all PG1A, PG1B, PG1C, PG2, PG7, PSG6, and CG5 (PSG1 and PSG2) isolates. Further to this, aminoglycoside resistance genotypes *APH(3’’)-Ib* was observed in all isolates with *APH(6)-Id* except CG5, and all PG1A, PG1B, and PG1C isolates possessed *APH(3’)-Ia*. Additional aminoglycoside resistance genes (*AAC(3)-IV* and *APH(4)-Ia*) were observed in PG1C and PSG6, *AAC(6’)-Ib7* was observed in all PG3 isolates, except isolate 39288, *aadA5* was observed in both PG6 isolates, and *ANT(3’’)-IIa* was observed in PSG2-4. The beta-lactamase gene *bla*_TEM-1_ and ESBL *bla*_CTX-M-15_ were present in all PG2 isolates together with the plasmid mediated quinolone resistance (PMQR) gene *qnrS*, *tet(A),* aminoglycoside resistance genes and *sul2*. The *qnrS* gene was also present with a range of different genes in the PG6 and PG7 isolates. The ESBL *TEM-150*, was observed in all PG6, PG7 and both CG5 (PSG2-3) isolates. Regarding trimethoprim resistance, four mutually exclusive genotypes were observed *via dfrA5, dfrA1, dfrA14,* and *dfrA17* where *dfrA5* was observed in all PG1A, PG1B, and PSG6 isolates, *dfrA1* was observed in all PG3 isolates (except isolate 39288) and in the PSG2-5 isolates, *dfrA14* was observed in both PG7 isolates, and *dfrA17* was observed in both PG6 isolates. Resistance to macrolides, lincosamides, and streptogramin B (*ermB*) was identified in all PG3 and PG6 isolates. Further macrolide resistance (*mphA*) was observed in all *ermB* positive PGs and in isolate 31 (PG7). Further phenicol resistance (*catB3* and *catI*) was observed in all PG3 isolates, except isolate 39288. Sulphonamide resistance was conferred *via* one of two mutually exclusive genes (*sul1* or *sul2*;) with *sul1* being observed in all PG3 isolates (except isolate 39288, which contained *sul1* on the chromosome), both PG6 isolates, and the PSG2-4 and PSG6 isolates. Comparatively, *sul1* was observed in all PG1A, PG2, PG7, and PSG6 isolates.

A total of 65 biocide resistance, 83 metal resistance genes, and 23 biocide and metal resistance genes were observed across the chromosomes of isolates sequenced for this study. Of these, 56 biocide resistance genes, 58 metal resistance genes, and 19 dual biocide-metal resistance genes were found to be ubiquitous (SI Table BACMET: Chromosome). From these, ubiquitous biocide resistance genotypes towards acetates, acids, acridines, alcohols, alkanes, cycloalkanes, imidazoles, organo-sulphates, organo-tins, paraquats, peroxides, phenanthridines, phenyls, quaternary ammonium compounds (QACs), thiaziniums, triarylmethanes, and xanthenes were observed. Additionally, ubiquitous metal resistance towards antimony, arsenic, bismuth, cadmium, chromium, cobalt, copper, iron, lead, magnesium, manganese, mercury, molybdenum, nickel, selenium, silver, tellurium, tungsten, vanadium, and zinc was observed. Further resistance (*via acrE* and *emmdR*) to acids, acridines, organo-sulphates, paraquats, phenanthridines, and QACs was observed in all isolates, except CG5. Further acid resistance was observed in all isolates except PG1C containing CG1 isolates (*evgA*); in all isolates except isolates 2 - 4 (CG3), isolate 5 (CG5) and both CG6 isolates (*gadA*); in all isolates except PG1B containing CG1 and CG4 (*ydeP*), in all non-CG1 isolates (*ymgB)* (associated with peroxide resistance); and in isolate 10 (CSG1) *(gadB)*. Further resistance to paraquats, phenanthridines, and QACs was observed in all CG4, CG7, CSG1, and CSG2 isolates *via emrE*. The biocide resistance gene *qacEΔ1* was observed in isolate 39288 (CG5). Further resistance to peroxides (*sitABC*) was observed in all PG1A and PG1B containing CG1 isolates, both CG5 isolates, and in CSG1-2 isolates. Finally, further acid and peroxide resistance (*yjaA*) was observed in all CG5 and CG7 isolates.

With regards to non-ubiquitous chromosomal metal resistance, all isolates except isolate 39286 (CSG3) contained *arsC*, an additional antimony and arsenic resistance gene, with isolate 10 possessing two more operonic arsenic-antimony resistance genes (*arsAD*) (SI Table BACMET Chromosome). All CG1-3, CG5, and CSG1 (isolate 10) isolates possessed further nickel and cobalt resistance *via fecDE*. Additional chromium resistance (*nfsA*) was observed in all non-CG1 isolates. Additional iron resistance was observed in all CG1 and CG5 isolates *via ybtPQ*. A large copper resistance operon (*pcoABCDERS*) and a large silver resistance operon (*silABCEFPRS*) was observed in all CG6 and CG7 isolates. The *G2alt* aluminium resistance gene was observed in both CG5 isolates. The *sitABC* operon observed in all PG1A and PG1B containing CG1 isolates, both CG5 isolates, and in CSG1-2 isolates is attributed to manganese and iron resistance. Finally, further cobalt resistance (*yjaA*) was observed in all CG5 and CG7 isolates.

Only one plasmid mediated biocide resistance gene was observed, *qacEΔ1*, conferring resistance to quaternary ammonium compounds. This gene was observed in all CG3 isolates (except isolate 39288), CG6, CG7, CSG1, and CSG2 isolates. While absent in the plasmid, *qacEΔ1* is present on the chromosome of isolate 39288, suggesting either a misclassification of a plasmid contig or a transposition event. Resistance to mercury was the only plasmid metal resistance operon detected, where the full *merACDEPRT* operon was observed in all PG1A, PG1B, PG3 and in isolates 34 and 10 (PSG4 and 6). Partial *mer* operons (*merACDEPT*) were observed in PG7, *merADE* was observed in both PG7 isolates, and *merDE* was observed in both CG5 (PSG1-2) isolates.

Between 74 and 111 virulence factor genes were observed in each chromosome (SI Table VFDB Chromosome). Of these, 48 were ubiquitously distributed which, in broad terms (as assigned by VFDB), confer the following predicted capabilities: adherence, biofilm, effector delivery system, immune modulation, motility, and nutritional or metabolic factor (in particular enterobactin siderophores). For adherence, all isolates except CG7 contained *ykgK/ecpR* and *fdeC*, all isolates except isolate 5 (CG5) and CG6 possessed *csgE*; all isolates except CG5 and CG7 possessed *fimAFGI*, *fimBE* were also absent from CG5 and CG7 and isolate 34 (CSG2) the *fimCDH* were present in one copy for all isolates and two copies in the majority of isolates; the *yagWVXYZ* adherence gene cluster was absent from CG3 and CG7, with just *yagW* being absent from CG5; the *papBCDEFHIJK* gene cluster was observed in all CG1 isolates with a partial cluster (*papCD*) also observed in isolate 34 (CSG2); the *afaABCDE-VIII* operon was observed in all CG1 isolates possessing PG1A; the *lpfABC* gene cluster was observed in all CG4 isolates and isolate 39286 (CSG3); finally, *sinH* was observed in both CG5 isolates and isolate 10 (CSG1). Regarding biofilm forming virulence factors, only one was observed (*algA*), which was contained in all CG1 bovine manure treated grass isolates (PG1A).

Regarding effector delivery systems, the *espLPRXY* (including its variants) were present in a mosaic distribution: all isolates except isolate 34 (CSG2) possessed *espX4*, *espX1* was also present in most *espX4*+ isolates with the exception of CG2, CG6, CG7, and isolate 10, one isolate (isolate 34) possessed *espX1* but not *espX4*; all isolates except CG5-7 possessed *espR1*; *espY1* was observed in CG5 and CG7 isolates and in isolate 10, isolate 10 also contained *espY2*, *espY3*, and *espY4*; *espL1* was observed in CG6 and isolate 10 while *espL4* was observed in CG5 and isolate 10; and *espP* was observed in the three CG1 isolates possessing PG1C. All isolates except CG6 possessed *hcp-2*; all isolates except CG1 isolates possessing PG1A, CG6, and CG7 contained *gspCDEFGHIJKLM*. Finally, *spaP* was observed in all CG-3, CG5-7, and isolates 10 and 39286 (CSG1 and CSG3). Only the *icsP/sopA* exoenzymes were detected in all CG1, CG4, isolates 39287 (CG5), 10, and 39286.

The immune modulation gene *rfaD* was observed in all CG1, CG5, and CG7 isolates whereas *rfaE* was observed in CG5-7, isolate 10, and isolate 39286; *manC* was observed in all CG3-4 isolates; both *fcl* and *gmd* were observed in all CG1 isolates possessing PG1C and CG5 isolates; finally *gtrA* was observed in isolate 34 only while *gtrB* was observed in both isolates 10 and 34. Regarding invasion factors, *aslA* was observed in all CG5-7 and isolate 10; *kpsDM* was observed in isolates 10 and 34, and *kpsT* was observed in isolate 10.

The motility gene, *flgI* was observed in all isolates except isolate 10 whereas *flgD* was observed in all CG1, CG5-7, and isolate 10. The *fliGMPQ* cluster was observed in all isolates except CG5 and all *fliGMPQ* isolates except CG7 also contained *fliI*. Finally, all PG1A isolates contained *papX*. Regarding siderophores, while enterobactin synthesis was ubiquitous, yersiniabactin, aerobactin, and salmochelin synthesis pathway proteins were observed. Yersiniabactin synthesis genes and clusters (*fyuA*, *irp1*, *irp2*, and *ybtAEPQSTUX*) were observed in isolate 34 and all CG1 and CG5 isolates. Aerobactin synthesis genes (*iutA* and *iucABCD*) were observed in isolate 10, all PG1B, and CG5. Salmochelin synthesis genes (*iroCDEN*) were only observed in isolate 39287 (CG5). Finally, the *chu* and *shu* heme utilisation clusters (*chuSUWY* and *shuATVX*) were observed in isolate 10 only. Regarding stress survival, only one gene (*clpP*) was observed in isolate 34, isolate 39286, and in CG2-CG7.

Plasmids in 28 isolates contained a virulence factor (SI Table VFDB: Plasmid). All PG1C isolates possessed the *faeCDEFHIJ* gene cluster enabling further adhesion capacity. The exoenzyme *icsP/sopA* was observed in all PG1A, PG1B, PG4, and PSG3-6 isolates. For siderophores, the salmochelin synthesis gene cluster *iroBCDEN* was observed in all PG1A, PG1B, PSG2, and PSG4 isolates. The aerobactin gene *iutA* and gene cluster *iucABCD* were observed in all PG1A isolates.

### Chromosomal point mutations associated with antimicrobial resistance

Chromosomal point mutations conferring resistance to nalidixic acid and ciprofloxacin were observed in 16, were phenotypically ciprofloxacin resistant. In addition isolate 10, which contained only one point mutation in the *parC* was susceptible (SI Table Pointfinder, Table 2). Mutations in the *gyrA* gene corresponding to amino acid 87 was observed in 16 isolates with all CG4 isolates with D87H and CG2, CG5, and isolate 34 with D87N. An additional *gyrA* mutation corresponding to the amino acid 83 (S83L) was observed in isolate 34 and all CG2, CG3, and CG5. A mutation conferring an amino acid change S80I in *parC* was observed in isolate 34, and all CG2 and CG5 isolates. Finally, a mutation conferring an amino acid change S57T in *parC* was detected in isolate 10.

## Discussion

Using a dual ecological and evolutionary approach for a One Health problem we aimed to shed light on antimicrobial resistance and potential pathogenicity in *E. coli* in agricultural manure and grassland. Isolates in this study were mainly of the O8:H25 or O8:H30 serogroups. While other serogroups and sequence types (including ST10) were observed, the prevalence and persistence of this serogroup was of interest from a One Health perspective, especially due to the relative stability of chromosomal gene content and differential plasmid gene content (CG1 and CG2). The presence of the O8 in CG1, CG2, and CG4 with a lack of an O antigen in CG3 and different O antigen in 39286 may suggest that O8 acquisition is ancestral to the divergence of CG1-4 from CG5-7. The absence of *E. coli* of the clinically promiscuous, resistant or multi-drug resistant and globally dominant ST131 demonstrates that these *E. coli* are not the same as the most frequently identified clinical *E. coli*. The porcine manure associated isolates were earlier diverging than bovine manure associated isolates, except for isolate 10, the earliest diverging isolate. The chromosomal proximity of CGs may offer an insight as to why certain chromosome groups (*e.g*., CG5 and CG7) repeatedly displayed the same genotypes as these were sister taxa. The pangenome of this study was relatively closed (with the sum of gene families remaining stable across the isolates) suggesting a relatively closely related subset of genomes^47^.

Tetracycline resistance was widespread in this study with 35 isolates phenotypically resistant, predominantly conferred by the *tetA* or *tetB* gene. The lack of these genes in other tetracycline resistant strains however suggests the possibility of an unscreened mutation or other mechanism. However, tetracycline resistance is predominantly mediated via tetracycline resistance genes. These results suggest that a bias towards tetracycline resistance exists in the agriculture as tetracycline was not used as a selective agent in the isolation of the bacteria. This could reflect previously observed per-weight antibiotic bias in Irish veterinary practice, with tetracyclines comprising 55.8%^48^.

All ciprofloxacin resistant isolates had point mutations in *gyrA* and *parC*. From a plasmid perspective, all isolates containing the *qnrS1* were bovine manure isolates, which were co-selected with an ESBL (*bla*_CTX-M-15_ or *bla*_TEM-150_) and tetracycline resistance gene. The *bla*_CTX-M-15_ positive isolates were only detected in bovine manure.

Resistance to kanamycin but not to amikacin was observed. Aminoglycosides were administered to bovines in the studies when required to treat infection and were not administered as a prophylactic. Aminoglycosides are recommended for use with caution in Irish veterinary practice and therefore are not the first line of treatment. As isolates from control soil and manure treated grass (within CG1) were also aminoglycoside resistant, the reservoir may be from the environment. From a genotype perspective, an array of aminoglycoside resistance efflux pumps was ubiquitously observed. The three CG1 plasmid groups all displayed a similar aminoglycoside resistance genotype (*APH(6)-Id, APH(3’’)-Ib,* and *APH(3’)-Ia*), with PG1C also possessing *AAC(3)-IV,* suggesting these genes to be aetiological of resistance. Interestingly, both *APH(6)-Id* and *APH(3’’)-Ib* were observed in PG2 which were also isolated from bovine manure treated grass but selected on cefotaxime or ciprofloxacin media. The control soil isolates also contained *APH(3’)-Ia*, which confers resistance to kanamycin but not amikacin. These results suggest that *APH(3’)-Ia* played the most active role in resistance and as the CG1 bovine manure treated grass isolates were extracted five weeks after initial spread, PG1A may have acquired this gene from PG1B isolates. As isolate 10 (bovine manure) also possessed *APH(3’)-Ia, APH(6)-Id* and *APH(3’’)-Ib* despite being an ancestral strain and possessing a differential plasmid group (PSG6) may further highlight this acquisition theory. The porcine manure isolates with kanamycin resistance all possessed *AAC(6’)-Ib7*, suggesting this to be the genetic mechanism of resistance in these isolates.

While zinc was incorporated into pig feed, no differential zinc resistance genotype was observed across isolates regardless of their source^2^. From a virulence genotype perspective, the presence of *icsP/sopA* across the phylogeny but particularly in CG1 (PG1A and PG1B) was of interest. This gene is associated with actin mediated intracellular locomotion in *Shigella* spp., which may suggest that these isolates may be enteroinvasive^49,50^. Further to this, the diverse siderophore production genes in CG1 compared to other isolates suggests that it may be adaptive to low iron environments (*e.g*., grass, most soils, or blood)^51^. From an evolutionary perspective, the chromosomal acquisition of *algA* (a *Pseudomonas* associated alginate-rich biofilm forming gene) on the chromosomes of all CG1 isolates from bovine manure treated grass was particularly interesting. To our knowledge, this is the first report of *algA* in *E. coli*, with *E. coli* most commonly forming biofilms using β-1,6-N-acetyl-D-glucosamine polymers, colanic acid, or cellulose^52^. As the human lung is a hostile environment subject to free radical DNA damage like what may be induced by UV, this gene may offer a survival strategy to cells on sun facing leaves^53,54^. Furthermore, *E. coli* presenting O8 and O17 are commonly associated with biofilm formation in clinical samples, suggesting a potential role in adaptability to diverse environments^55,56^. Finally, the presence of *astA* (a heat stable enterotoxin) within isolate 34 three weeks after application suggests that toxicogenic strains may survive in exposed environments but as this was only observed in one isolate, more work is required to determine the validity of this observation.

These results suggest that *E. coli* in soils and grasses may adapt to their new environments evolving novel strategies. Due to the bias towards biofilm forming virulence factors and the relatively large abundance of O-antigen types associated with increased biofilm formation, manure spreading may facilitate introducing persistent and potentially pathogenic bacteria to grassland environments. Depending on how long the antimicrobial resistance genes survive, they may be reintroduced to grazing animals at a later stage, potentially weakening the effect of prescribed antibiotics, and posing a One Health threat. Greater analysis of persistence of these bacteria and research into their transmission from primary elements of the food chain are required to understand their risk to animal and human health.

## Author contributions

CT: Investigation, Writing – Original Draft, Methodology, Data Curation; CMB: Project Administration, Resources, Supervision; FPB: Supervision, Project Administration, Resources; DM: Investigation, Data Curation; DD: Data Curation, Resources, Supervision; RJL: Data Curation, Writing – Original Draft, Formal Analysis, Methodology, Visualization; FW: Conceptualization, Funding Acquisition, Project Administration, Validation, Writing – Review & Editing, Supervising

## Acknowledgements

This project was funded by a Walsh Fellowship (2017037) to Dr Ciara Tyrrell. We thank the staff who contributed and managed the field trial in the Teagasc field centre in Wexford.

